# High fat high sucrose exposure during lactation but not during the post-weaning period programs impaired glucose homeostasis

**DOI:** 10.1101/2025.01.15.633246

**Authors:** Anne Goergen, Marcela Borsigova, Stephan Wueest, Daniel Konrad

## Abstract

Nutrition during critical developmental windows plays a pivotal role in shaping long-term metabolic health. Exposure to a Western diet in early life may lead to altered metabolic programing, potentially influencing the risk to develop obesity and type 2 diabetes. In particular, an early exposure to an obesogenic diet may induce a metabolic program that only emerges upon dietary challenges later in life. Accordingly, understanding the long-term impact of nutritional insults in different stages of early life may provide strategies for primary prevention. Therefore, the present study aimed to determine which time window in the post-natal period is most critical for long-term metabolic health. To this end, pups born to lean mouse dams were exposed to a high-fat, high-sucrose (HFHS) diet for 3 weeks during lactation or in the post-weaning period. Thereafter, mice were either fed a regular chow diet until the age of 30 weeks or exposed to a second bout of HFHS diet for the last 12 weeks. Metabolic health and development of obesity was assessed by regular monitoring of glucose homeostasis and body weight gain. 12 week-old mice with lactational exposure to a HFHS diet revealed a significantly impaired glucose tolerance compared to control mice, while glucose tolerance of post -weaning exposed mice was similar to the control group. Moreover, a second bout of HFHS diet impaired glucose tolerance and increased body and adipose tissue weight in mice with lactational exposure to a significant greater extent than in post-weaning exposed mice. Hence, exposure to HFHS during lactation but not during the post-weaning period programs impaired glucose homeostasis, suggesting that the lactational window may be more critical in terms of metabolic programing.

## 1. Introduction

The globally rising prevalence of obesity in children and adolescences is one of the biggest public health challenges of the 21^st^ century (WHO). Childhood obesity is a strong predictor for the development of adult obesity, which is strongly associated with cardio-metabolic disturbances such as insulin resistance and type 2 diabetes (T2D), metabolic-associated fatty liver disease (MAFLD), cardiovascular disease (CVD) and cancer, eventually leading to premature death (Singh et al. 2008; Simmonds et al. 2015; Twig et al. 2016; Li et al. 2016; Zheng, Ley, and Hu 2018; Vucenik and Stains 2012; Whitlock et al. 2009). It becomes increasingly apparent that genetic predisposition, a sedentary lifestyle, and/or overnutrition alone cannot explain the exponential increase in obesity observed over the last four decades (Dyer and Rosenfeld 2011). A substantial body of evidence suggest that nutritional imbalance at early stages of life may favor obesity and metabolic dysfunction in adulthood (Butruille et al. 2019; Lukaszewski et al. 2013; Bouret, Levin, and Ozanne 2015; Vogt et al. 2014). For example, obese children were shown to display elevated biomarkers of inflammation, which has been linked to cardiovascular disease in later life (Skinner et al. 2010). The idea that a nutritional stimuli or insult, experienced during critical stages of development, permanently alters an individual’s physiology and metabolism is defined as metabolic programming (Dyer and Rosenfeld 2011). The exposure to such events may happen during various life stages. Hence, fetal programming describes effects of *in utero* events, while the effects during weaning (suckling) are referred to as lactational or post -natal programming (Cerf 2015). This concept is supported by human epidemiological studies which demonstrate that breastfed neonates were protected from rapid weight gain and increased susceptibility to obesity when compared to formula fed infants (Owen et al. 2005). In line, numerous rodent studies have shown that early and time-limited exposure to overnutrition may lead to deterioration of glucose metabolism and increased adiposity in adulthood (Masuyama and Hiramatsu 2014; Tsuduki et al. 2013; Glavas et al. 2010; Glavas et al. 2021; Liang et al. 2016). In particular, it was shown that exposure to a high-fat diet (HFD) confined to the lactation period is particularly detrimental in respect to metabolic health of the offspring since it affects leptin sensitivity and adipose tissue expansion (Sun et al. 2012b; Butruille et al. 2019). Moreover, exposure to HFD during suckling impairs neuronal innervation of hypothalamic neurocircuits critical for glucose homeostasis and decreases parasympathetic outputs to pancreatic β-cells, thereby causing disturbances in glucose and energy metabolism (Vogt et al. 2014). Collectively, these studies support the notion that lactation is a critical time window for metabolic programing which can affect glucose and energy homeostasis as well as adiposity later in life.

Besides, the post-weaning period may be considered a critical developmental window for long-term metabolic health. In rodents, transitioning onto regular chow diet upon weaning leads to a marked decline in dietary fat intake, from 20-45 % fat content in maternal milk to ∼ 11.5 % fat in chow diet (Görs et al. 2009). Similarly, in humans, the transition f rom maternal milk to solid food represents a crucial phase in life, which is typically accompanied by a shift in the lipid content of the diet: f rom lipid-rich breast-milk to a higher carbohydrate content in solid food (Shaoul, Tiosano, and Hochberg 2016). However, introduction of solid food upon weaning in infants may often expose them to diets richer in fat than recommended, since fat enriched diet is cheap and readily available (Meldrum, Morris, and Gambone 2017; WHO 2018).

Developmental programming paradigms often include a “second -hit”, in which a second exposure to a specific (metabolic) insult after the initial programming event leads to the manifestation of an aggravated metabolic phenotype. In humans, such a second-hit may be the prolonged exposure to a Western diet during adulthood. Given the pervasive exposure to obesogenic diets in many societies, a Western diet trigger is a clinically relevant stressor for humans who have experienced metabolic programming insults in early life. While several studies have reported adult phenotypes in rodents exposed to “Western diets” during lactation, little is known about adult metabolic phenotypes of rodents exposed to metabolic insults in the post-weaning period (Vogt et al. 2014; Butruille et al. 2019; Monks et al. 2018). In particular, comparison of the long-term impact of an exposure to a Western diet during the lactational and the post - weaning period remains to be unraveled.

Since interventional studies investigating different nutritional paradigms in humans, and especially children, are considered difficult and may even be unethical, we used a rodent model to investigate the impact of early life exposure to a high-fat, high-sucrose (HFHS) diet, representative of the human Western diet. In particular, we aimed to determine the long-term susceptibility to obesity and glucose intolerance resulting f rom HFHS exposure in the post-weaning compared to the lactational period. In addition, we aimed to investigate whether a second exposure to a HFHS diet in previously exposed mice leads to an aggravated metabolic phenotype.

## 2. Methods

### 2.1 Animal husbandry and experiments

All animal experiments were conformed to the Swiss animal protection laws and were approved by the Cantonal Veterinary Office in Zurich, Switzerland (License ZH293/2018). C57BL6/J mice were purchased from Charles River (Sulfzeld, Germany). All experiments were performed with mice kept in a 12h: 12h light: dark cycle (light phase starting at 7 a.m.) in a pathogen-free animal facility. Mice were housed 2-5 mice per cage, in individually ventilated cages. The ambient temperature in the animal facility was kept constant at 22 °C and the animals were fed standard rodent (chow) diet or HFHS diet (43 % fat, 40 % sucrose, ssniff-Spezialdiäten GmbH, Soest, Germany), with *ad libitum* access to food and water.

### 2.2 Physiological tests

For intraperitoneal insulin tolerance tests (ipITT), animals were fasted at 9 a.m. After 3h of fasting, animals were weighed and a small incision into the tail was made using a razor blade for blood sampling. Plasma glucose was measured before injection of insulin (Actrapid® HM Penfill, Novo Nordisk Pharma AG) and 15, 30, 45, 60, 90 and 120 min after injection. After measuring baseline glucose levels, mice were injected intraperitoneally with 0.75 or 1 U/kg body weight human recombinant insulin. Blood glucose levels were measured with a blood glucometer (Accu-Check Aviva, Roche Diagnostics).

For intraperitoneal glucose tolerance tests (ipGTT), animals were fasted at 5 p.m. until 8 a.m. the next morning. After overnight fasting, animals were weighed and a small incision in the tail was made using a razor blade for blood sampling. Plasma glucose was measured before injection of glucose and 15, 30, 45, 60, 90 and 120 min after injection. After measuring baseline glucose levels, the mice were injected intraperitoneally with 2 mg/kg body weight glucose (Glucosum “Bichsel” 20%, Grosse Apotheke Dr. G. Bichsel AG, Switzerland). Blood glucose levels were measured with a blood glucometer. In order to assess glucose stimulated insulin secretion, 50 μl of blood were sampled from a small incision in the tail before intraperitoneal glucose injection as well as 10 and 30 minutes after glucose injection. Blood was processed as described below.

### 2.3 Blood and tissue harvest

Before euthanasia, animals were fasted for 5 hours (8 a.m. to 1 p.m.). Blood glucose was measured immediately before euthanasia as described above. Each mouse was euthanized individually using CO_2_ and death was determined by the absence of withdrawal reflexes of the lower limbs. The abdominal cavity was opened immediately after euthanasia, and systemic blood was sampled after cardiac puncture using 30G syringes. Blood was transferred to an Eppendorf tube containing 5 mmol/L ethylenediaminetetraacetic (EDTA, pH 8) as anticoagulant. After centrifugation at 8000 rpm at 4 °C for 10 min, plasma was stored at −80 °C until further processing. Adipose tissue depots, skeletal muscle (quadriceps) and liver were carefully dissected, weighed on an analytical scale and snap frozen in liquid nitrogen. Tissue samples for RNA isolation were snap frozen in RNA Tissue protect reagent (Qiagen) before snap freezing. Popliteal lymph nodes were removed from inguinal adipose depots. Plasma and tissue samples were kept at -80 °C until further processing.

### 2.4 Insulin ELISA

Plasma insulin levels were measured using the kit Ultra Sensitive Mouse Insulin ELISA Kit (Chrystal Chem, Catalogue # 90080, Zaandam, Netherlands).

### 2.5 Statistics and software

All results are expressed as mean ± standard error of the mean (SEM). Statistical analysis was performed using two-tailed, unpaired Student’s *t*-test, one-way ANOVA or two-way ANOVA with Tukey’s multiple comparisons (GraphPad Prism Software, San Diego, CA, USA; Version 8.0.0), assuming normal distribution. Outliers were defined as mean+-2 * standard deviations (SD) and were excluded from statistical analysis. For α=5, ^#^p<0.1, *p<0.05, **p< 0.01 and ***p< 0.001, ****p<0.0001. GraphPad prism was used to produce graphs and figures.

## 3. Results

### 3.1 Impaired glucose tolerance of lactation and post-weaning HFHS exposed mice at 6 weeks of age

Nutrition during critical windows of development plays a pivotal role in shaping long-term metabolic health. In order to understand what time window of metabolic insults mainly affect metabolic fate in mice, we defined two 3-week nutritional windows, in which young mice were subjected to early HFHS exposure. These early exposure corresponded to HFHS exposure during lactation (indirect consumption via dam’s milk, from birth until weaning at postnatal day 21, defined as “lactation exposed mice”, lactation HFHS-chow) or exposure to HFHS after weaning (direct consumption, from postnatal day 21 to postnatal day 42, defined as “post-weaning exposed mice”, post-weaning HFHS-chow). Mice fed a chow diet throughout their first 6 weeks of life were used as control mice (defined as “control mice”, chow-chow). At the age of 6 and 12 weeks, glucose tolerance and/or insulin sensitivity was assessed in all groups. To investigate whether a second exposure to a HFHS diet in previously exposed mice lead to an aggravated metabolic phenotype, we exposed a subset of cohorts of each group to a second bout of HFHS at the age 18 weeks until the end of the experiment when mice were 30 weeks old. Before terminating the experiment, glucose tolerance and insulin sensitivity was assessed in all mice. Early exposed mice fed a chow diet from 18-30 weeks of life were defined as chow-fed control mice, chow-fed lactation exposed mice and chow-fed post-weaning exposed mice.

Mice exposed to a second bout of HFHS were defined as HFHS-diet challenged control mice (Chow-HFHS), HFHS-diet challenged lactation exposed mice (Lactation HFHS-HFHS) and HFHS-diet challenged post-weaning exposed mice (post-weaning HFHS-HFHS)(Figure 1).

**Figure 1.**
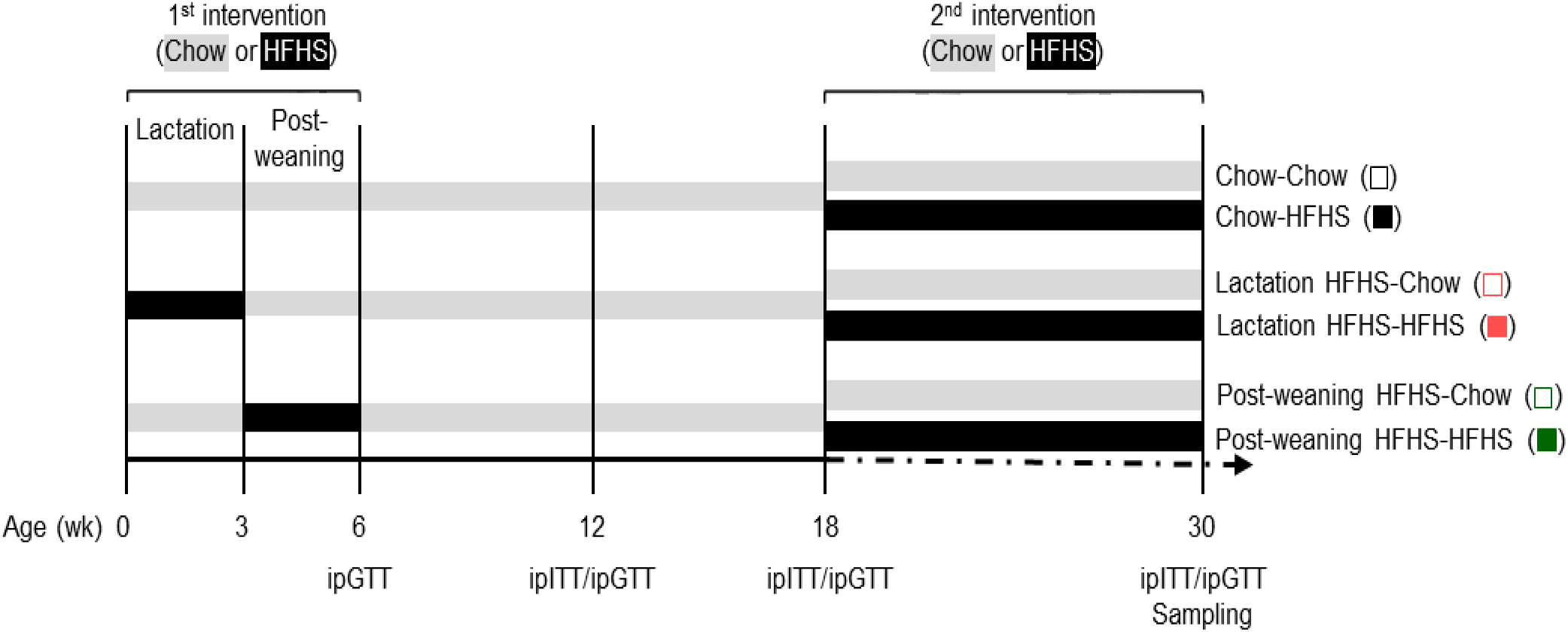
Experimental design and time points of experimental interventions. Chow-Chow: Chow-fed control mice. Chow-HFHS: HFHS diet challenged control mice; Lactation HFHS-Chow: Chow-fed lactation exposed mice. Lactation HFHS-HFHS: HFHS diet challenged lactation exposed mice. Post-weaning HFHS-Chow: Chow-fed post-weaning exposed mice; Post-weaning HFHS-HFHS: HFHS diet challenged post-weaning exposed mice. Wk: weeks. ipGTT: glucose tolerance test. ipITT: insulin tolerance test.

At 6 weeks of age, both lactation exposed and post-weaning exposed male mice displayed significantly impaired glucose tolerance in comparison to control mice (Figure 2A). This finding is reflected by a significantly higher area under the curve (AUC) of the intraperitoneal glucose tolerance test (ipGTT) of both early exposed groups compared to control mice (Figure 2B). Similarly, lactation exposed and post-weaning exposed female mice had significantly impaired glucose tolerance 15 minutes after glucose injection (Figure 2C) leading to a significantly increase in the AUC in post-weaning exposed mice (Figure 2D). Taken together, exposure to HFHS diet during the lactation and the post-weaning time period leads to impaired glucose tolerance in male and female mice at 6 weeks of age.

**Figure 2:**
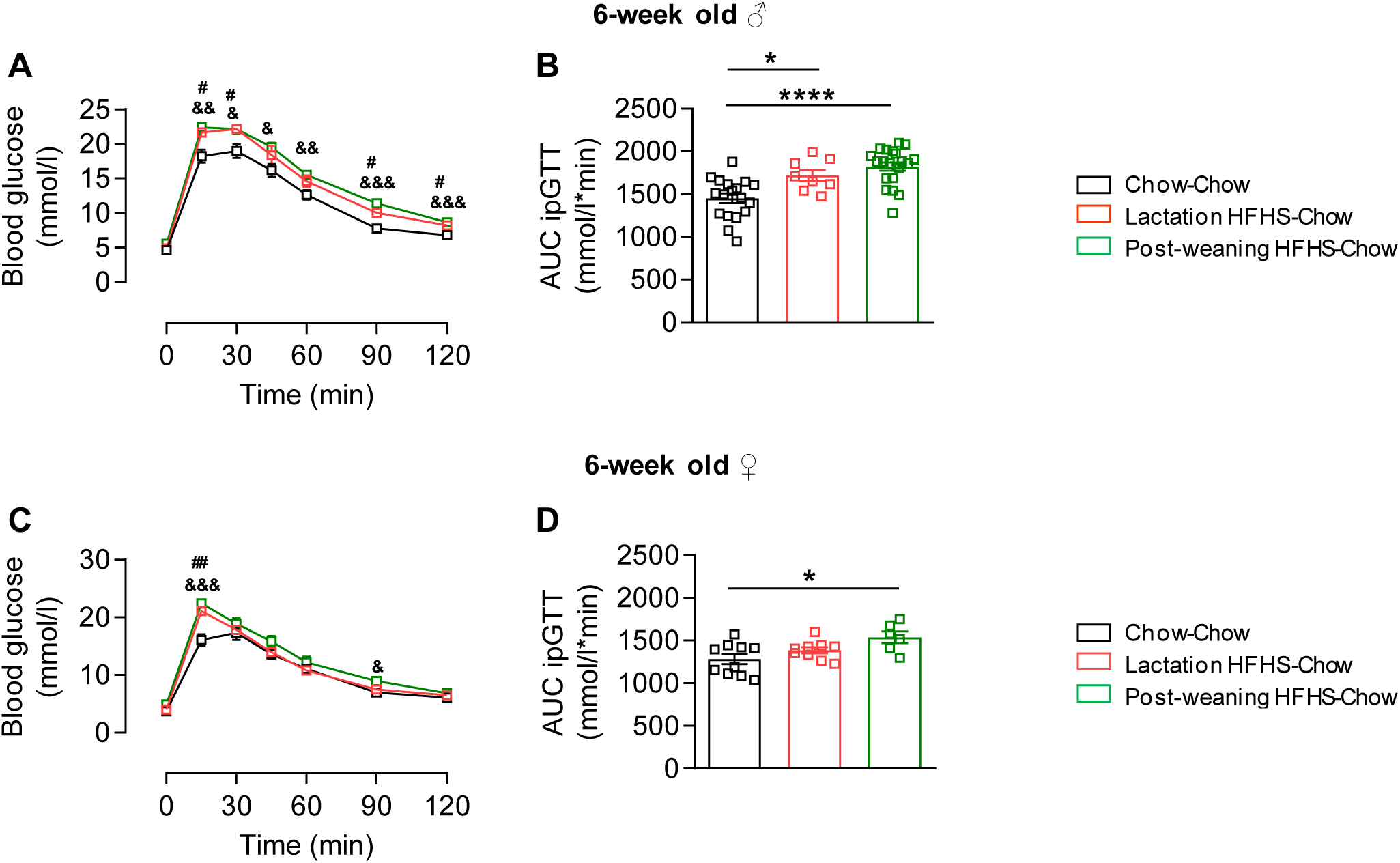
Impaired glucose tolerance of lactation and post-weaning HFHS exposed mice at 6 weeks of age. **A.** Intraperitoneal glucose tolerance test (ipGTT) in 6 weeks-old male mice (n=8-21). **B.** Area under the curve (AUC) of blood glucose levels during ipGTT of male mice at 6 weeks of age (n=8-21). **C.** ipGTT in 6 weeks-old female mice (n=6-11). **D.** AUC of blood glucose levels of female mice at 6 weeks of age (n=8-21). Data are shown as mean ± SEM. *p<0.05, **p<0.01, ***p<0.001 ****p<0.0001. (One-way ANOVA for panels B and D; Two-way ANOVA for panels A and C, Post hoc analysis was performed using Tukey’s multiple comparisons test). Symbols represent statistical differences between groups in panels A and C: & - Chow-Chow vs. Post-weaning HFHS-Chow. # - Chow-Chow vs. Lactation HFHS-Chow.

At 6 weeks of age, both lactation exposed and post-weaning exposed male mice displayed significantly impaired glucose tolerance in comparison to control mice (Figure 2A). This finding is reflected by a significantly higher area under the curve (AUC) of the intraperitoneal glucose tolerance test (ipGTT) of both early exposed groups compared to control mice (Figure 2B). Similarly, lactation exposed and post-weaning exposed female mice had significantly impaired glucose tolerance 15 minutes after glucose injection (Figure 2C) leading to a significantly increase in the AUC in post-weaning exposed mice (Figure 2D). Taken together, exposure to HFHS diet during the lactation and the post-weaning time period leads to impaired glucose tolerance in male and female mice at 6 weeks of age.

### 3.2 Impaired glucose tolerance in lactation HFHS exposed mice at 12 weeks of age

In order to understand the long-term impact of HFHS exposure within important developmental time windows on whole-body metabolism, we assessed glucose and insulin tolerance at 12 weeks of age. Glucose intolerance persisted in lactation exposed male mice, while it was no longer different between post-weaning exposed and control male mice (Figure 3A). In line, AUC of the ipGTT was significantly higher in lactation exposed male mice, both in comparison to chow-fed and post-weaning exposed age-matched male mice (Figure 3B). Additionally, glucose-stimulated insulin secretion was higher in pre-weaning exposed mice, indicating a compensatory increase in insulin secretion to handle the glucose bolus (Figure 3C). In contrast, insulin sensitivity was similar in all groups in 12 weeks-old male mice, as shown by similar AUC of intraperitoneal insulin tolerance test (ipITT) (Figure 3D). These data suggest that impaired glucose tolerance in lactation exposed mice originates from a β-cell defect rather than from decreased insulin sensitivity. Similarly, females exposed to HFHS during lactation depicted impaired glucose tolerance at the age of 12 weeks, as suggested by significantly higher plasma glucose levels during ipGTT and higher AUC in comparison to control mice (Figure 3E, F). In contrast to male mice, glucose tolerance in post-weaning exposed female mice was still slightly impaired at 12 weeks of age (Figure 3E, F). Of note, the AUC after glucose injection was significantly higher in lactation exposed mice than in post-weaning exposed mice at 12 weeks of age (Figure 3F). Moreover, both groups showed impaired ipITT compared to chow-fed controls albeit the difference between post-weaning exposed and control females was only trend wise increased) (Figure 3G). Collectively, lactation exposed male and female mice displayed impaired glucose metabolism at 12 weeks of age, while glucose tolerance in post-weaning exposed mice was only slight impaired in females but normalized in males.

**Figure 3:**
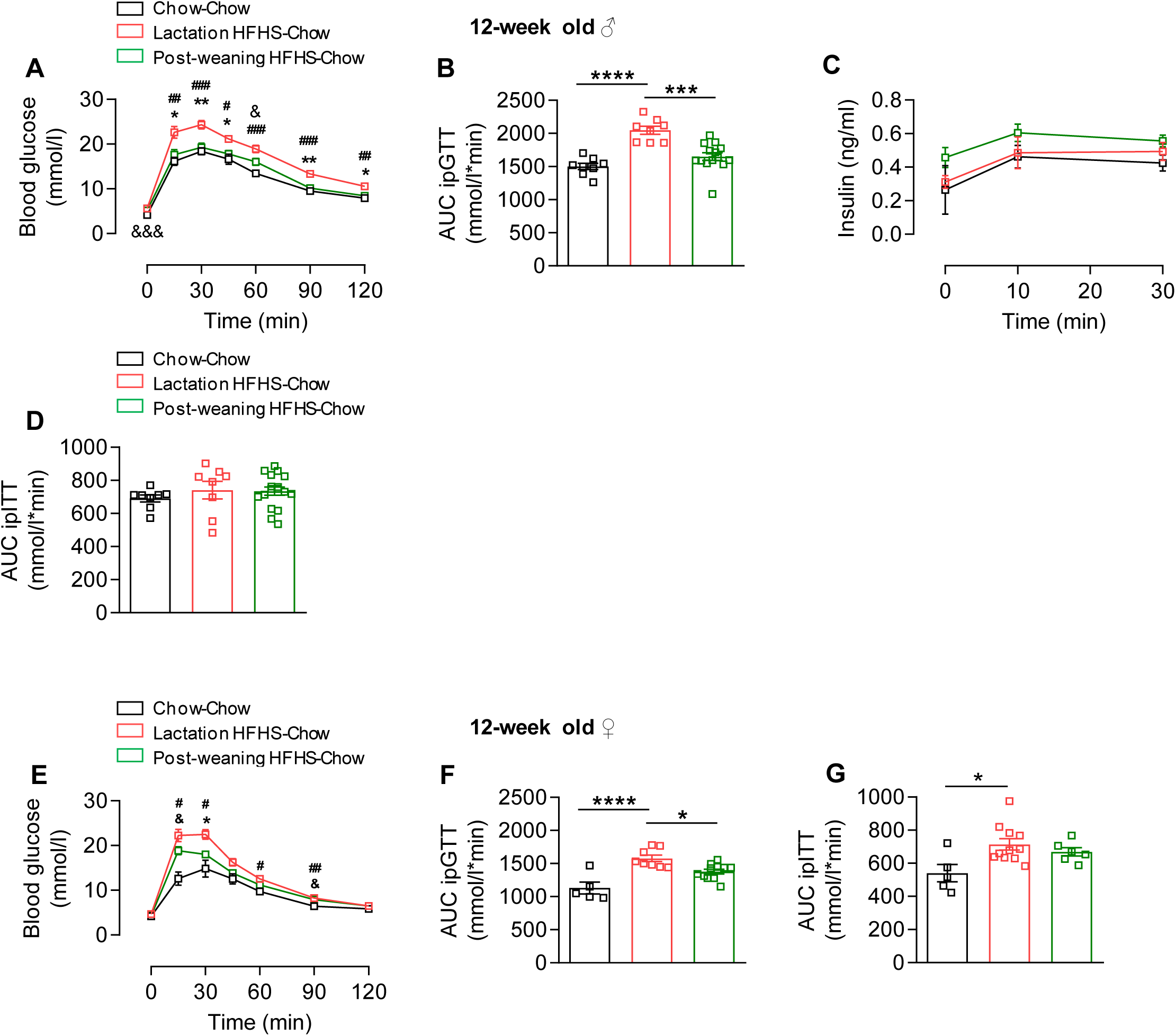
Impaired glucose tolerance in lactation HFHS exposed mice at 12 weeks of age. **A.** ipGTT in 12 weeks-old male mice (n=8-21). **B.** AUC of blood glucose levels during ipGTT (n=8-21). **C.** Insulin levels (ng/ml) during ipGTT (n=5-6). **D.** AUC of blood glucose levels during ipITT (n=8-21). **E.** ipGTT in 12 weeks-old female mice (n=6-11). **F.** AUC during ipGTT (n=8-21). **G.** AUC during ipITT (0.75 U/kg) n=5-11). Data are shown as mean ± SEM. *p<0.05, **p<0.01, ***p<0.001, ****p<0.0001. (One-way ANOVA for panels B, D, F and G; Two-way ANOVA for panels A, C and E, Post hoc analysis was performed using Tukey’s multiple comparisons test). Symbols represent statistical differences between groups in panels A and E: & - Chow-Chow vs. Post-weaning HFHS-Chow. # - Chow-Chow vs. Lactation HFHS-Chow. * - Lactation HFHS-Chow vs. Post- weaning HFHS-Chow

### 3.3 Detoriated glucose tolerance in HFHS diet challenged lactation HFHS exposed male mice at 30 weeks of age

In order to determine the effect of a second exposure to a HFHS diet on metabolic health in pre-exposed mice, we challenged a subset of control, lactation and post-weaning exposed 18 week-old mice with a second bout of HFHS diet for 12 weeks. At the age of 30 weeks, there was no difference in glucose tolerance between male chow-fed control, lactation exposed and post-weaning exposed mice (Figure 4A). In mice challenged with a second exposure to HFHS diet, basal glucose levels were significantly higher after overnight fasting in lactation exposed mice, both in comparison to control and post-weaning exposed mice. Moreover, ipGTT was significantly impaired in HFHS diet challenged lactation exposed mice in comparison to HFHS-diet challenged control and post-weaning exposed mice (Figure 4B). Accordingly, the AUC of HFHS diet challenged lactation exposed males was significantly higher when compared to the AUC of HFHS diet challenged control and post-weaning exposed mice (Figure 4C). Chow-fed post-weaning exposed mice had blunted insulin sensitivity in comparison to chow-fed lactation exposed males, as suggested by significantly higher glucose levels during ipITT (Figure 4D). In contrast, insulin sensitivity was similar in all three groups challenged with a second bout of HFHS diet (Figures 4E and 4F). Collectively, our data suggest that while glucose tolerance is similar in chow-fed control, lactation and post-weaning exposed 30 weeks-old male mice, a second exposure to HFHS diet is more detrimental for glucose homeostasis in lactation than post-weaning exposed male mice. Conversely, insulin sensitivity is impaired in chow-fed post-weaning exposed 30 week-old males in comparison to age-matched lactation exposed mice, while a second exposure to HFHS diet does not reveal a significant difference in insulin sensitivity between both early exposed groups suggesting that early exposure to HFHS rather primes the metabolic organs for glucose intolerance than for insulin resistance in male mice.

**Figure 4:**
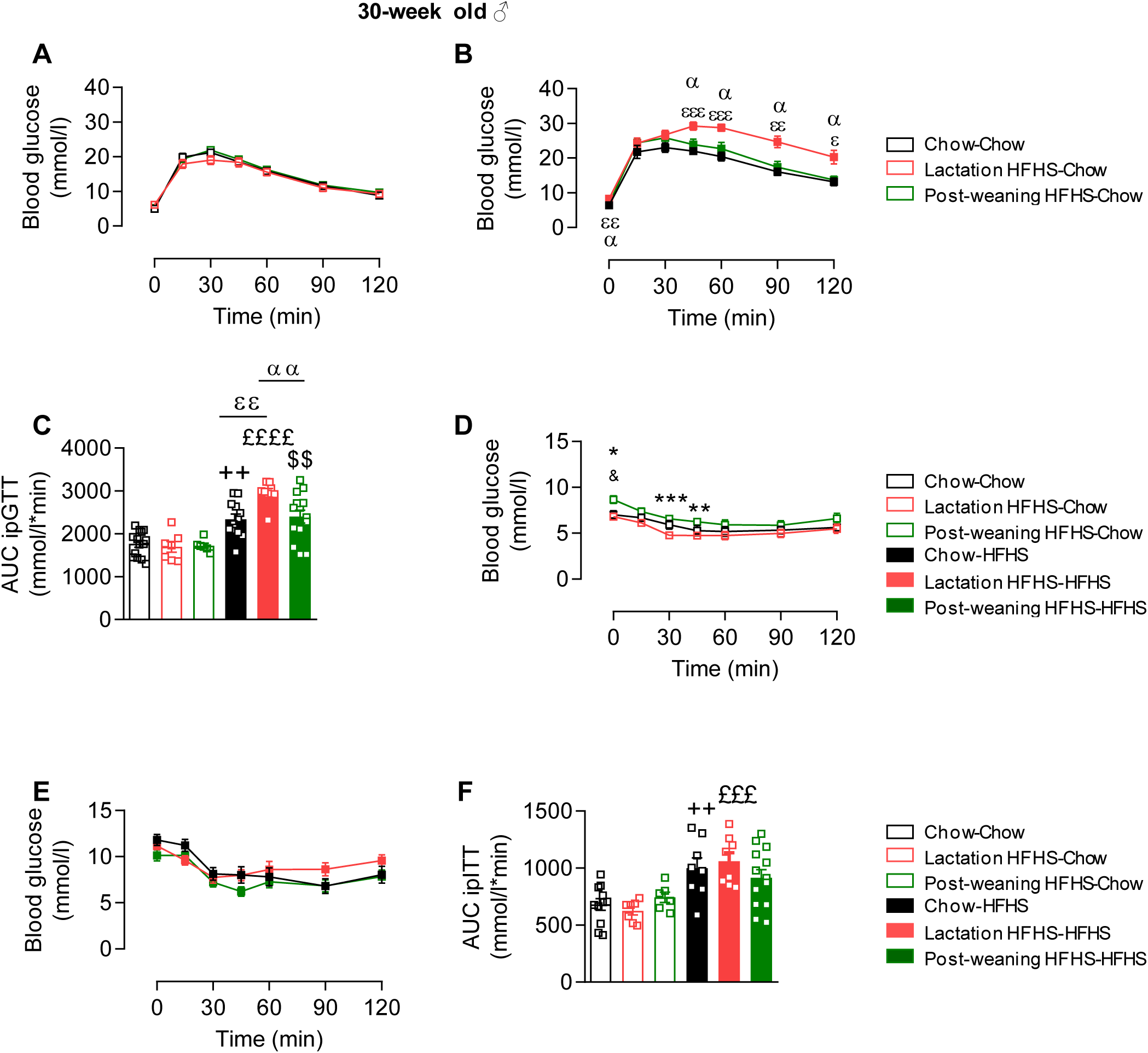
Glucose tolerance of chow-fed and HFHS diet challenged lactation and post-weaning HFHS exposed male mice at 30 weeks of age. **A.** ipGTT in 30 weeks-old chow-fed male mice (n=7-14). **B.** ipGTT in 30 weeks-old HFHS diet challenged male mice (n=8-14). **C.** AUC during ipGTT in 30-weeks old chow-fed male and HFHS-diet challenged male mice (n=8-14). **D.** ipITT (1 U /kg) in 30 weeks-old chow-fed male mice (n=7-14). **E.** ipITT (1 U /kg) in 30 weeks-old HFHS-challenged male mice (n=7-14). **F.** AUC during ipITT in chow-fed male and HFHS-diet challenged 30 weeks-old male experimental mice (n=7-14). Data are shown as mean ± SEM. *p<0.05, **p<0.01., ***p<0.001. (Two-way ANOVA for all panels; Post hoc analysis was performed using Tukey’s multiple comparisons test) Symbols represent statistical differences between groups: & - Chow-Chow vs. Post-weaning HFHS-Chow, + - Chow-Chow vs. Chow- HFHS, £ - Lactation HFHS-Chow vs Lactation HFHS-HFHS, $ - Post-weaning HFHS-chow vs Post-weaning HFHS-HFHS, α - Lactation HFHS-HFHS vs. Post-weaning HFHS-HFHS, δ - Chow-HFHS vs. Post-weaning HFHS-HFHS. ε - Chow-HFHS vs. Lactation HFHS-HFHS

### 3.4 Detoriated glucose tolerance in HFHS diet challenged lactation and post-weaning HFHS exposed female mice at 30 weeks of age

When assessing glucose tolerance in 30 week-old females that were not exposed to a second bout of HFHS, we found that glucose tolerance of lactation exposed females was significantly impaired compared to control females (Figure 5A). In females receiving a second bout of HFHS diet, only lactation exposed female mice displayed a significantly higher AUC during ipGTT in comparison to the corresponding chow-fed control group (Figure 5B, C). Insulin sensitivity was significantly reduced in lactation exposed females in comparison to chow-fed control and post-weaning exposed female mice (Figure 5D). Of note, only HFHS diet challenged lactation exposed female mice displayed a significantly higher AUC after glucose injection in comparison to the corresponding chow-fed control group (Figure 5D). Furthermore, insulin sensitivity was similar in all groups exposed to a second bout of HFHS (Figure 5E, F). Taken together, a second bout of HFHS diet only significantly impaired glucose tolerance in lactation exposed females, while chow-fed lactation exposed females displayed impaired insulin sensitivity in comparison to control females suggesting that exposure to HFHS during lactation severely affects glucose metabolism in C57Bl6/J female mice.

**Figure 5:**
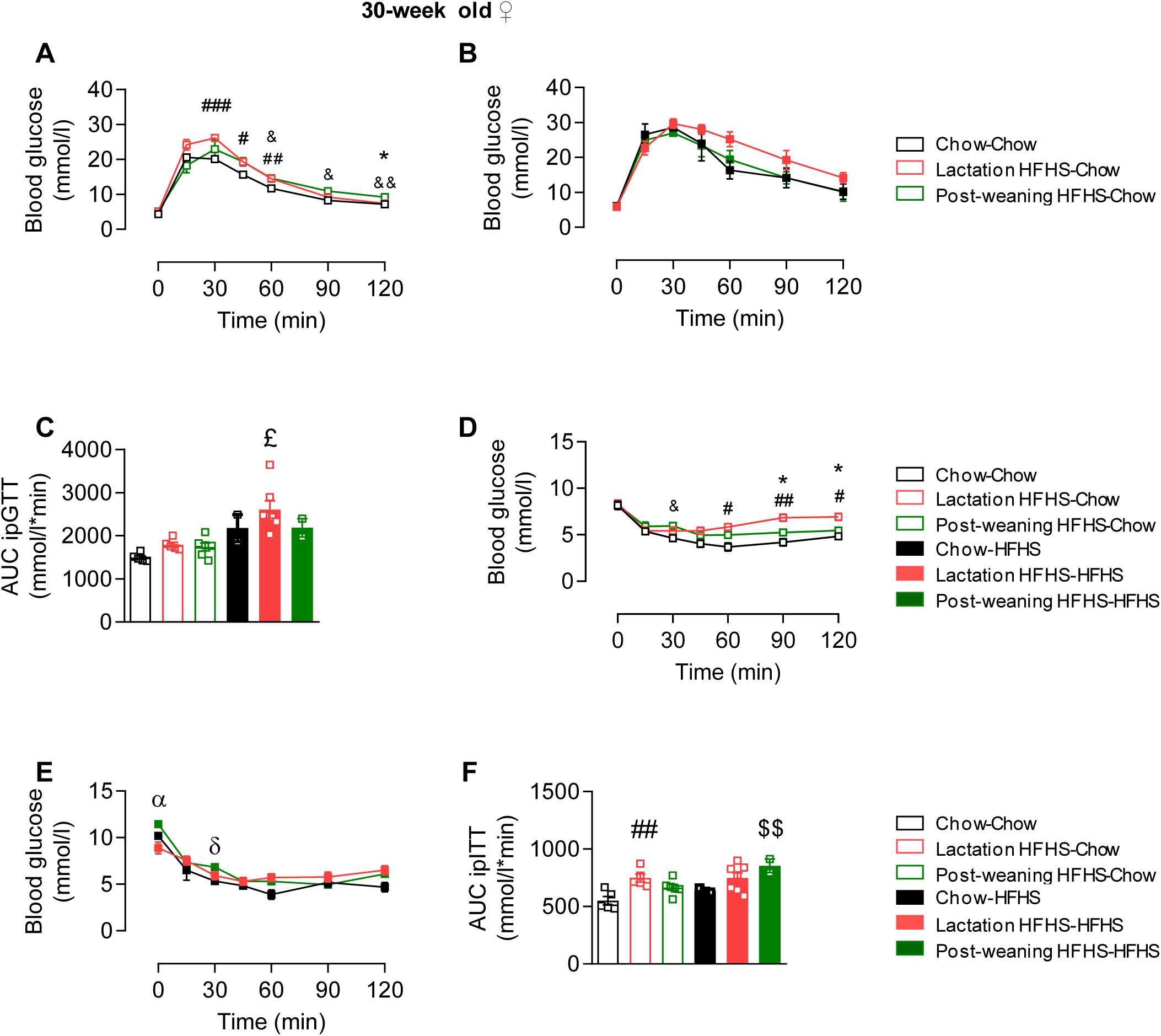
Detoriated Glucose tolerance in HFHS diet challenged lactation HFHS exposed female mice at 30 weeks of age. **A.** ipGTT (2 mg /kg) in 30 weeks-old chow-fed female mice (n=5-8). **B**. ipGTT in 30 weeks-old HFHS diet challenged female mice (n=5-8). **C.** AUC during ipGTT (n=5-8). **D.** ipITT (1 U kg) in 30 weeks-old chow-fed female mice (n=5-8). **E.** ipITT in 30 weeks-old HFHS diet challenged female mice (n=5-8) **F.** AUC during ipITT (n=5-8). Data are shown as mean ± SEM. *p<0.05, **p<0.01., ***p<0.001. (Two-way ANOVA for all panels; Post hoc analysis was performed using Tukey’s multiple comparisons test). Symbols represent statistical differences between groups: & - Chow-Chow vs. Post-weaning HFHS-Chow, # - Chow-Chow vs. Lactation HFHS-Chow, * - Lactation HFHS-Chow vs Post-weaning HFHS-chow, + - Chow-Chow vs. Chow-HFHS, £ - Lactation HFHS-Chow vs Lactation HFHS-HFHS, $ - Post-weaning HFHS-chow vs Post-weaning HFHS-HFHS α - Lactation HFHS-HFHS vs. Post-weaning HFHS-HFHS, δ - Chow- HFHS vs. Post-weaning HFHS-HFHS.

### 3.5 Body and organ weight of chow-fed and HFHS diet challenged lactation and post-weaning HFHS exposed male mice

Since increased body and fat mass may negatively affect glucose homeostasis, these parameters were assessed next. There was no body weight difference between chow-fed control, lactation and post-weaning exposed male mice (Figure 6A, B). In line, fat pad and liver weights were comparable between the three groups (Figures 6C, D, E, F, G, H). As expected, a second exposure to a HFHS diet led to a significant increase in body weight gain, a significantly increased final body weight, as well as significantly higher fat pad and liver weights in all three groups compared to age-matched chow-fed males. Moreover, HFHS diet challenged lactation exposed mice gained significantly more weight than HFHS diet challenged post-weaning exposed mice, paralleled by significantly increased white and brown adipose tissue as well as liver weight (Figure 6A-H).

**Figure 6:**
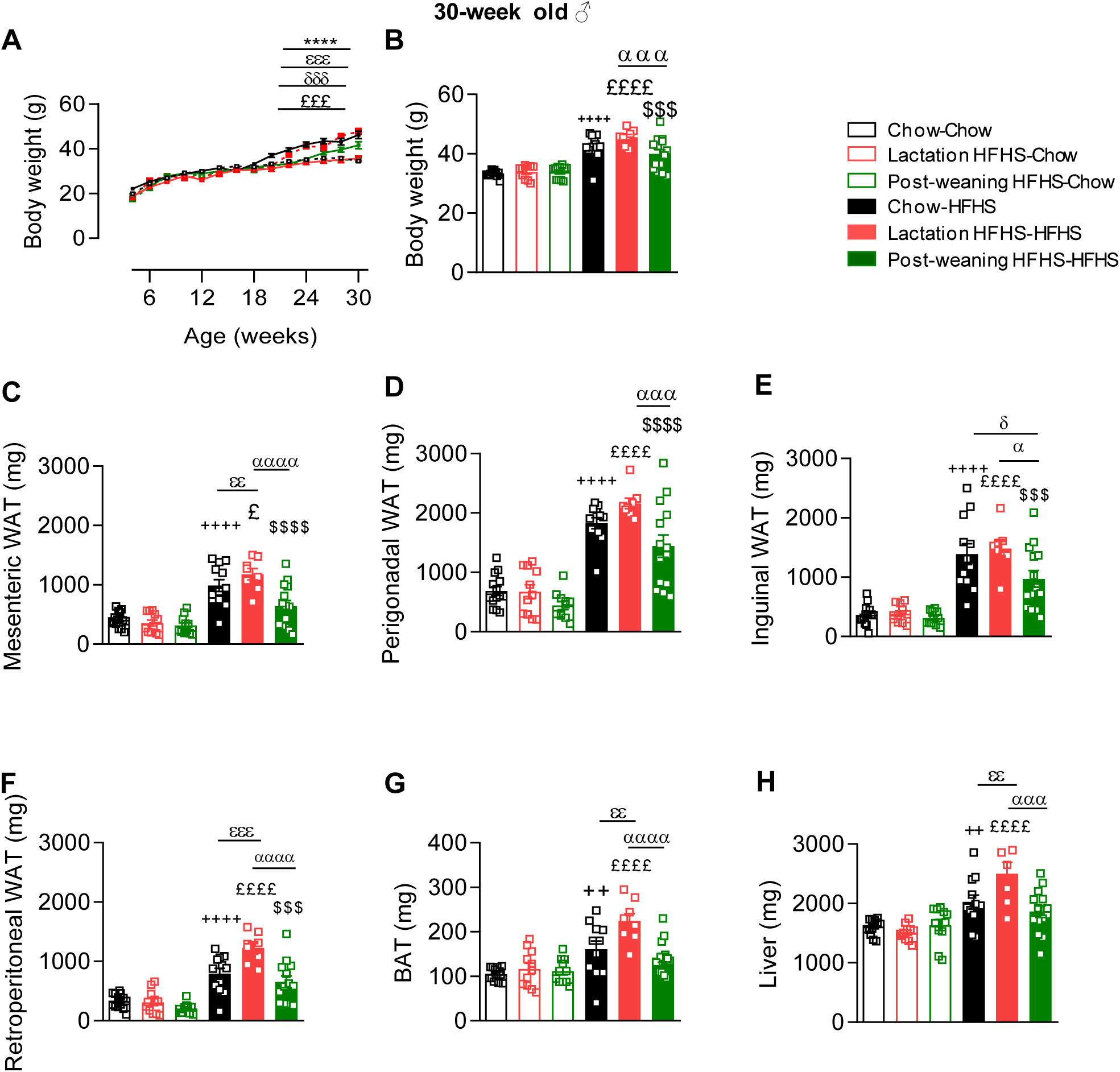
Body and organ weight of chow-fed and HFHS diet challenged lactation and post-weaning HFHS exposed male mice. Body weight development (grams) over time (weeks) in male mice (n=6-14). **B.** Final body weight in male mice at the age of 30 weeks (n=10-14). Weight (mg) of mesenteric (**C**), perigonadal (**D**), inguinal (**E**) and retroperitoneal (**F**) adipose tissue of 30 weeks-old male mice (n=8-14). Weight (mg) of brown adipose tissue (BAT; **G**) and liver (**H**) of 30 weeks-old male mice (n=8-14). Data are shown as mean ± SEM. *p<0.05, **p<0.01., ***p<0.001, ****p<0.0001. (Two-way ANOVA for all panels, Post hoc analysis was performed using Tukey’s multiple comparisons test). Symbols represent statistical differences between groups: & - Chow-Chow vs. Post-weaning HFHS-Chow, # - Chow-Chow vs. Lactation HFHS-Chow, * - Lactation HFHS-Chow vs Post-weaning HFHS-chow, + - Chow-Chow vs. Chow-HFHS, £ - Lactation HFHS-Chow vs Lactation HFHS-HFHS, $ - Post-weaning HFHS-Chow vs Post-weaning HFHS-HFHS, α - Lactation HFHS-HFHS vs. Post-weaning HFHS-HFHS, δ - Chow-HFHS vs. Post-weaning HFHS-HFHS. ε - Chow-HFHS vs. Lactation HFHS-HFHS.

### 3.6 Body and organ weight of chow-fed and HFHS diet challenged lactation and post-weaning HFHS exposed female mice

Similarly, there was no difference in body, fat pad or liver weight between chow-fed control, lactation and post-weaning exposed females (Figure 7A-H). Intriguingly, liver weight was significantly higher in HFHS diet challenged lactation exposed compared to HFHS diet challenged post-weaning exposed females. Of note, only lactation exposed and post-weaning exposed females challenged with a second bout of HFHS diet were significantly heavier than age-matched chow-fed females, which was paralleled by significantly increased perigonadal, brown adipose tissue and liver weight (7A, B, D, G, H).

**Figure 7:**
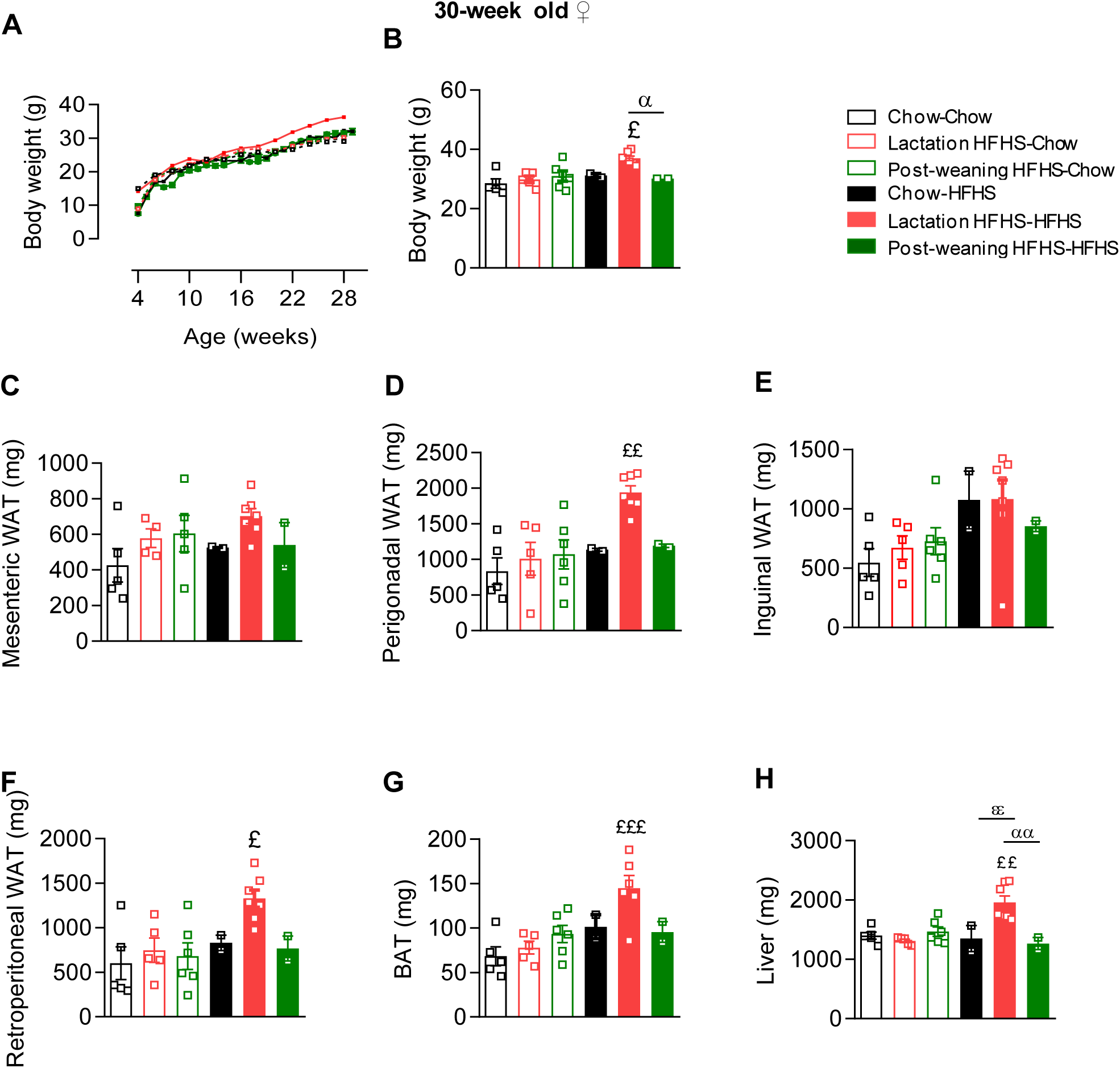
Body and organ weight of chow-fed and HFHS diet challenged lactation and post-weaning HFHS exposed female mice. **A.** Body weight development (grams) over time (weeks) in female mice (n=2-7). **B.** Final body weight in female mice at the age of 30 weeks (n=2-7). Weight (mg) of mesenteric (**C**), perigonadal (**D**), inguinal (**E**) and retroperitoneal (**F**) white adipose tissue of 30 weeks-old female mice (n=2-7). Weight (mg) of brown adipose tissue (BAT; **G**) and of liver (**H**) in 30 weeks-old female mice (n=2-7). Data are shown as mean ± SEM. *p<0.05, **p<0.01., ***p<0.001. (Two-way ANOVA for all panels, Posthoc analysis was performed using Tukey’s multiple comparisons test). Symbols represent statistical differences between groups: £ - Lactation HFHS-Chow vs Lactation HFHS- HFHS, α - Lactation HFHS-HFHS vs. Post-weaning HFHS-HFHS, δ - Chow-HFHS vs. Post-weaning HFHS-HFHS.

## 4. Discussion

Early exposure to an obesogenic diet may induce a metabolic program that only emerges upon further dietary challenges later in life. Accordingly, understanding the long-term impact of nutritional insults in different stages of early life may provide strategies for primary prevention. For this reason, the present study aimed to determine which time window in the post-natal period is most critical for long-term metabolic health. Regular monitoring of glucose homeostasis and body weight gain revealed that the timing of HFHS diet exposure during critical development periods differentially primes glucose metabolism and weight gain in mice. In particular, we demonstrate that glucose tolerance in chow-fed lactation exposed mice remained impaired as long as 9 (males and females) or 27 (females) weeks after the initial metabolic insult. In mice exposed to HFHS diet after weaning, glucose tolerance was significantly impaired in males and females directly after the intervention, but approached glucose tolerance of chow-fed control mice thereafter. However, post-weaning exposure to HFHS diet induced impaired insulin sensitivity in male mice at 30 weeks of age. A second bout of HFHS diet significantly impaired glucose tolerance and increased body as well as adipose tissue weight in mice with lactational compared to post-weaning exposure. Hence, exposure to HFHS during lactation but not during the post-weaning period programs impaired long-term glucose intolerance and highlights sex-dependent differences in metabolic programing related to insulin sensitivity.

We compared mice exposed to a 3-week HFHS feeding period during two important development periods: the lactation period (from postnatal day 0 to postnatal day 21) and the post-weaning period (from postnatal day 21 to postnatal day 42). Both time frames are associated with periods of rapid body weight gain as well as developmental mile stones, such as maturation of the hypothalamo-pituitary-gonadal axis (Holtrup et al. 2017; Ahima, Prabakaran, and Flier 1998). During growth spurts, metabolic organs may be particularly sensitive to stressors, such that developmental insults potentia**ll**y alter metabolic outcomes and may manifest as obesity, insulin resistance, and type 2 diabetes (Devaskar and Thamotharan 2007). In this respect, it is not surprising that we found impaired glucose tolerance at 6 weeks of age in both groups exposed to an early HFSH diet and it indicates that metabolic insults during important growth spurts are detrimental for glucose metabolism. Of note, glucose tolerance was similarly impaired in post-weaning and in lactation exposed mice at 6 weeks of age, even though lactation exposed mice were fed a regular chow-diet for 3 weeks before the ipGTT.

Further evidence for a more detrimental impact of a HFHS diet exposure during lactation is provided by the finding that at 12 weeks of age, i.e. 9 weeks after the initial metabolic insult, lactation exposed males still displayed deteriorated glucose tolerance, while post-weaning exposed male mice, with a chow-fed period of only 6 weeks, had a glucose tolerance that was comparable to age-matched control mice. Moreover, we observed that post-weaning but not lactation exposed males demonstrated a compensatory increase in glucose-stimulated insulin levels compared to control mice, likely resulting in significantly impaired glucose tolerance in lactation but not post-weaning exposed mice. In line, Vogt et al previously reported that exposing mice to HFD during lactation leads to a deterioration in glucose homeostasis by decreasing parasympathetic innervation of pancreatic β-cell islets, causing decreased insulin output after glucose stimulation (Vogt et al. 2014). Hence, impaired glucose tolerance in lactation exposed mice likely originated from β-cell damage.

Similar to 12-week old males, lactation exposed female mice had significantly worsened glucose tolerance compared to post-weaning exposed mice. However, in contrast to males, there was an additional deterioration in insulin sensitivity in lactation exposed females, suggesting that such impairment contributed to the deteriorated glucose tolerance in these mice.

Of note, the finding that lactation exposed mice have impaired glucose tolerance at 12 weeks of age is opposed to findings by Glavas et al. who report that lactation exposed Swiss Webster mice at 10 weeks of age don’t have impaired glucose tolerance compared to mice who received chow diet during suckling (Glavas et al. 2021). We speculate that this discrepancy may be due to strain-related differences in obesity and diabetes susceptibility.

With regard to long-term metabolic health in chow-fed early exposed mice, we found that glucose tolerance was no longer different between lactation and post-weaning exposed male mice at 30 weeks of age. However, it was intriguing to observe that post-weaning exposed males demonstrated impaired insulin sensitivity. We speculate that the relative hyperinsulinemia observed in 12-week old post-weaning exposed mice may have led to a decline in insulin sensitivity with aging, in line with studies demonstrating that hyperinsulinemia induces insulin resistance and/or type 2 diabetes (Kolb et al. 2020; Marín-Juez et al. 2014; Nolan et al. 2015). Interestingly, the phenotypic difference in insulin sensitivity between lactation exposed male and female mice observed at 12 weeks of age remained in 30 weeks old mice, since insulin sensitivity was still impaired in lactation exposed females. Furthermore, also glucose intolerance persisted in lactation exposed females, indicating that exposure to HFHS diet during lactation affected female mice more severely.

Several studies have previously shown that metabolic insults, especially during suckling, increase the susceptibility to develop obesity and associated metabolic comorbidities (Hafner et al. 2019; Vogt et al. 2014; Monks et al. 2018; Butruille et al. 2019). However, much less is known about the effects of metabolic insults in the post-weaning time period on long-term susceptibility to metabolic disorders. Hence, we compared metabolic parameters of HFHS diet-challenged lactation and post-weaning exposed mice at 30 weeks of age to investigate which time period of early exposure is most detrimental for the metabolic phenotype when challenged with a second metabolic insult. HFHS diet challenged lactation exposed male and female mice both displayed impaired glucose tolerance and increased body weight compared to post-weaning exposed mice. In parallel, adipose tissue and liver weights were increased in male and female mice exposed to HFHS diet during lactation. Thus, our study suggests that nutritional stimuli during lactation are more critical for metabolic programing than the dietary environment in the post-weaning period, independent of sex.

Even though the post-weaning phase is an important developmental time period with significant increases in body mass and adipose tissue development, we speculate that lactation is even more important because many organs maturate in mice during suckling (Holtrup et al. 2017; Ellsworth et al. 2018). For instance, metabolically relevant tissues such as skeletal myocytes, β-cells and hepatocytes complete maturation during suckling in rodents, while murine adipose tissue development extends until several weeks post-weaning and, thus, remains sensitive to developmental cues after lactation (Ellsworth et al. 2018).

In our study, both female and male mice were included, because the existence of sex -specific differences in metabolism is increasingly recognized in biomedical research (Dearden, Bouret, and Ozanne 2018). However, we generally saw consistent results between sexes, namely that glucose homeostasis is most affected in lactation exposed mice. This finding is opposed to results reported by several other studies. For instance, Glavas et al. reported that females were protected from hyperglycemia regardless of length or timing of HFD exposure (Glavas et al. 2021). Moreover, a study by Sun et al. demonstrated that males but not females exposed to a HFD during lactation led to decreased leptin sensitivity in the hypothalamus and thus impaired energy homeostasis (Sun et al. 2012a). One sex-specific difference that we observed was that chow-fed lactation exposed females but not males did develop insulin resistance. As an increasing number of studies start to examine the impact of maternal diet in both sexes, there is compelling evidence demonstrating that females may be more sensitive to exposure to increased glucose levels during early life. A likely candidate to mediate this effect may be estradiol. For example, it was recently shown that exposure to estradiol in early life periods induces hypothalamic insulin resistance in male rodents (Wang et al. 2018). Moreover, human studies have shown that female offspring is more prone to metabolic programing by diabetic parents than men (Krishnaveni et al. 2010; Gerlini et al. 1986). We therefore speculate that insulin sensitivity in lactation exposed female mice may have been impaired to a greater extent than in males by the HFHS. Finally, these discrepancies highlight the importance of including males and females in metabolic studies, since programming during the developmental periods may differentia**ll**y affect metabolic fate sex-dependently.

